# Carbon source-dependent metabolic states govern redox homeostasis and cofactor biosynthesis in *Propionibacterium freudenreichii*

**DOI:** 10.64898/2026.06.08.730882

**Authors:** Kirsi Savijoki, Bhawani Chamlagain, Minnamari Edelmann, Kaisa Hiippala, Paulina Deptula, Susanna Kariluoto, Tuula A. Nyman, Vieno Piironen, Pekka Varmanen

**Affiliations:** Department of Food and Nutrition, University of Helsinki, P.O. Box 66, FI-00014 University of Helsinki, Finland; Faculty of Medicine, University of Helsinki, P.O. Box 21, FI-00014, University of Helsinki, Finland; Department of Food Science, University of Copenhagen, Rolighedsvej 26, 1958 Frederiksberg, Denmark (PDeptula); Faculty of Pharmacy, Division of Pharmaceutical Chemistry and Technology, University of Helsinki, P.O. Box 56, FI-00014 University of Helsinki, Finland; Institute of Clinical Medicine, Department of Immunology, Faculty of Medicine, Rikshospitalet, University of Oslo and Oslo University Hospital, Norway (TA Nyman)

**Keywords:** Propionibacterium, proteomics, carbon metabolism, amino acid biosynthesis, vitamin B12, cellular redox status, organic acids, oxidative stress

## Abstract

Microbial adaptation to fluctuating nutrient and oxygen conditions requires coordinated regulation of metabolic networks to maintain redox homeostasis within physicochemical and energetic constraints. While oxygen-dependent responses in *Propionibacterium freudenreichii* (*PFR*) have been characterized at the transcriptomic level, the role of carbon source in defining system-level metabolic states remains unclear. Here, we investigated carbon source-dependent metabolic reprogramming and cofactor biosynthesis in *PFR* strain DSM 20271ᵀ using label-free quantitative proteomics integrated with physiological and metabolite analyses. Distinct carbon sources defined discrete metabolic states shaped by redox balance and flux distribution. Lactate supported a comparatively balanced physiological state characterized by enhanced respiratory metabolism, amino acid biosynthesis, and riboflavin metabolism, enabling high specific vitamin B12 yields (∼100 µg g^−1^ wet biomass). In contrast, hexose metabolism (glucose and fructose) imposed a redox-constrained state marked by upregulation of transport systems, glycolysis, and the pentose phosphate pathway, resulting in increased biomass but reduced biosynthetic efficiency. A defining feature of the hexose-driven state was activation of aspartate metabolism. Proteomic and metabolite data, together with functional assays, support a model in which aspartate is converted to fumarate and subsequently reduced to succinate, providing an alternative electron sink that facilitates NADH reoxidation under redox-constrained conditions. Together, these findings establish that carbon source shapes physiological state through flux distribution, redox homeostasis, and resource allocation, with cofactor biosynthesis emerging as a system-level property rather than a simple consequence of biosynthetic enzyme abundance.

**IMPORTANCE:** *Propionibacterium freudenreichii* is a central bacterium used in food fermentations and one of the few microorganisms able to synthesize biologically active vitamin B12, making it valuable for industry and biotechnology. Yet the metabolic principles that govern its performance under different growth conditions remain poorly understood. Here, we show that carbon source is a key determinant of metabolic state, dictating how cells resolve redox constraints and allocate biosynthetic resources. We uncover a previously unrecognized adaptation in sugar-grown cells, where aspartate functions as an alternative electron sink to sustain redox balance under constrained conditions. By contrast, lactate supports a physiological state that promotes efficient vitamin B12 biosynthesis. These findings reveal a central role for carbon source in shaping metabolic configuration and identify redox balancing as a critical lever linking environmental inputs to biosynthetic output. More broadly, this work provides mechanistic insight into redox-constrained metabolism and a framework for improving vitamin B12 production and other microbial bioprocesses.

## INTRODUCTION

*Propionibacterium freudenreichii* (PFR) is a metabolically versatile actinobacterium adapted to fluctuating anaerobic and microaerobic environments, where maintenance of redox homeostasis is central to growth and biosynthesis [1,2]. Its metabolism is organized around the Wood-Werkman cycle, which couples carbon flux from lactate and other substrates to propionate formation, ATP generation, and NADH reoxidation [3]. More broadly, metabolic output in PFR is limited by the interplay between carbon flux distribution, redox homesostasis, and energy conservation, consistent with constraint-based models describing how physicochemical, thermodynamic, and stoichiometric limits shape metabolic fluxes [4–6]. Carbon source availability strongly influences this organization, leading to substrate-dependent physiological states that reflect both genomic variability and regulatory flexibility [7,8].

PFR is widely used in food fermentation and is one of the few microorganisms capable of synthesizing bioactive vitamin B12 (hereafter B12). Because plants generally do not contain B12, microbial production of the vitamin is important for human nutrition, particularly in plant-based diets [9,10]. Beyond its applied relevance, this species provides a useful model for investigating how physiological behavior emerges from interacting environmental and metabolic constraints. PFR can utilize a broad range of substrates, including carbohydrates and organic acids such as lactate [1,7,8]. During anaerobic growth on lactate, carbon is channelled through the Wood–Werkman cycle, yielding propionate, acetate, and succinate. This pathway includes B12-dependent reactions, most notably methylmalonyl-CoA mutase, directly linking cofactor availability to flux distribution [1,11]. Changes in substrate utilization are therefore expected to influence both redox homeostasis and cofactor biosynthesis. Vitamin B12 biosynthesis in PFR proceeds through a complex de novo pathway encoded by approximately 30 genes [12]. The pathway begins with uroporphyrinogen III and culminates in assembly of the corrin ring, which contains a central cobalt ion and distinct upper and lower ligands [13,14]. The lower ligand, 5,6-dimethylbenzimidazole (DMBI), is synthesized from riboflavin in an oxygen-dependent reaction, creating a requirement for both anaerobic and microaerophilic conditions during B12 production [15,16]. This hybrid pathway creates an inherent mismatch: conditions that favor B12-dependent metabolism do not necessarily coincide with those that maximize biosynthetic capacity [17].

Oxygen adds a further layer of complexity by requiring coordination between respiratory and fermentative metabolism to maintain redox homeostasis [18,19]. In PFR, oxygen influences energy metabolism, stress adaptation, respiration, and cofactor biosynthesis [19–21]. Transcriptomic and surfaceome analyses have revealed extensive remodeling of transport systems and stress-response pathways during oxygen exposure [21]. At the same time, emerging omics studies indicate that nutrient composition affects energy generation, precursor availability, and B12 production [22,23]. Notably, B12-dependent metabolism becomes increasingly important under low-oxygen conditions despite reduced biosynthetic capacity, revealing a mismatch between cofactor demand and synthesis [17]. Together, these observations suggest that carbon source and oxygen jointly determine cellular physiology through a multilayered regulatory architecture integrating carbon metabolism, redox homeostasis, and cofactor biosynthesis [24].

Here, we investigated how carbon source influences cellular physiology, redox homeostasis, and cofactor biosynthesis in PFR strain DSM 20271ᵀ. Using label-free quantitative proteomics integrated with physiological and metabolite analyses, we compared cells grown on lactate, glucose, and fructose following an initial screening of five substrates. This approach revealed coordinated changes in transport systems, central carbon metabolism, amino acid pathways, and B12 biosynthesis. We further identified an aspartate-dependent fumarate-to-succinate pathway as a potential alternative electron sink supporting redox homeostasis during hexose metabolism. Together, these findings establish a systems-level framework linking substrate utilization to physiological adaptation and reveal broader principles of redox-constrained bacterial metabolism.

## MATERIALS AND METHODS

### PFR strain and culture conditions

The genome-sequenced PFR type strain DSM 20271 [25] was obtained from DSMZ (Braunschweig, Germany). The strain was routinely maintained on YEL agar (1% [w/v] yeast extract, 0.5% [w/v] tryptone, and 0.8% [w/v] sodium DL-lactate) under anaerobic conditions at 30 °C for 5-6 days and subcultured two to three times in liquid YEL medium (3-4 days per subculture). Prior to inoculation, cells were harvested, washed with 0.9% (w/v) NaCl, resuspended in an equal volume of 0.9% NaCl, and diluted 100-fold into YE medium (1% [w/v] yeast extract, 0.5% [w/v] tryptone, and 0.1 M K₂HPO₄/KH₂PO₄ buffer, pH 6.4) supplemented with 0.8% (w/v) sodium DL-lactate (LAC; Sigma-Aldrich), D-glucose (GLC; Merck), D-fructose (FRC; Merck), lactose (LTS; Buchs, Switzerland), or inositol (INO; Sigma-Aldrich). Carbon sources were supplied at identical mass concentrations (0.8%, w/v) to facilitate comparison of substrate-specific physiological responses; carbon-equivalent normalization was not applied. To prevent Maillard reactions, media without carbohydrates were autoclaved separately (121 °C, 15 min) and substrates were added aseptically. When indicated, cobalt chloride hexahydrate (5 mg/L; Sigma-Aldrich) was added to support B12 synthesis.

Colony forming ability of PFR cells in response to selected carbon sources (LAC, GLC, and FRC) was studied as follows. Cells were first cultured in YE medium supplemented with the indicated carbon source under the conditions described above. After four days (LAC- and GLC-grown cultures) and six days (FRC-grown cultures), cells were serially diluted (10⁻⁴ to 10⁻⁸) in phosphate-buffered saline (Dulbecco A, pH 7.3; Thermo Fisher Scientific). Five microliters of each dilution were spotted onto YE agar containing 0.8% (w/v) LAC, GLC, or FRC. Plates were incubated at 30 °C under anaerobic (AnaeroGen™ 2.5 L sachets and anaerobic jar; <1% O₂, ∼10% CO₂), semiaerobic (∼20% O₂, 5% CO₂), or aerobic (21% O₂, <0.1% CO₂) conditions for 4, 6, and 6-8 days, respectively. To assess the effect of aspartate on colony formation, cells were first cultivated in LAC-supplemented YE broth, serially diluted, and spotted onto YE agar containing LAC or FRC with or without 10 mM aspartate. Plates were incubated under the same atmospheric conditions as described above.

### Growth rate determination

Pre-cultures grown for 48 h were diluted 1:100 into fresh, unbuffered YE medium supplemented with the indicated carbon sources (0.8% [w/v]). Aliquots (300 µL) were dispensed into Bioscreen C honeycomb plates (four biological replicates) and incubated at 30 °C for 144 h in a Bioscreen C system (Oy Growth Curves Ab Ltd., Turku, Finland). Optical density (OD_600_) was measured every 4 h after 10 s shaking. Specific growth rates (h⁻¹) were calculated from mid-exponential phase OD values.

### Cultivation for metabolite and vitamin analyses

Cells were cultivated in 20 mL YE medium supplemented with LAC, GLC, or FRC (three biological replicates). Control cultures without cells and without added carbon sources were included. Cultures were incubated anaerobically at 30 °C for 72 h, followed by 96 h under microaerophilic conditions (150 rpm shaking). Microaerophilic conditions were generated by briefly opening and closing tube caps under sterile conditions. Growth (OD₆₀₀) and pH were measured at 72 h and 168 h. Cells were harvested at 168 h by centrifugation (12,000 × g, 10 min), washed with phosphate-buffered saline (PBS; pH 7.3), and stored at -20 °C together with supernatants for subsequent analyses.

#### Vitamin B12 analysis

Vitamin B12 analysis with a UHPLC system was based on the method described previously [26]. Cell pellets (0.05-0.3 g) were extracted in 10 mL acetate–NaOH buffer (pH 4.5) containing 1 mg sodium cyanide and incubated at 100 °C for 30 min. After cooling, samples were centrifuged twice (6,900 × g, 10 min), adjusted to 25 mL, filtered (0.2 µm), and analyzed using a Waters UHPLC system with photodiode array detection (361 nm) and an HSS T3 C18 column. Gradient elution (0.32 mL/min) was performed using water/acetonitrile (5-40%) containing 0.025% (v/v) trifluoroacetic acid. Vitamin B12 was quantified using a six-point external calibration curve of cyanocobalamin (Supelco, Bellefonte, PA, USA). Results were reported as volumetric (µg mL^−1^) and biomass-specific (µg g^−1^ fresh weight) B12 yields.

#### Riboflavin

Riboflavin (vitamin B2) levels in culture supernatants were determined by UHPLC with fluorescence detection as described previously [27]. Samples were acid-extracted, enzymatically treated (Taka-diastase and β-amylase), filtered (0.2 µm), and analyzed using a BEH C18 column with excitation/emission at 432/520 nm. Riboflavin was quantified based on an external standard curve of riboflavin (0.01– 1.0 ng µL-1).

#### Determination of sugars and organic acids

Sugars and organic acids were determined by a HPLC method as described previously [28]. Samples were diluted (1:10), filtered (0.45 µm), and a chromatographic separation was performed using a Hi-Plex H column (Agilent, CA, USA; 300 x 6.5 mm) and 10 mM H₂SO₄ as a mobile phase (0.6 mL/min, 65 °C). Acids were detected at 210 nm (UV), and sugars by refractive index detection. External calibration curves (0.1-1 mg mL^−1^) were used for quantification.

#### Amino acid analysis

Free amino acids were determined on an Acquity UPLC system according to the manufacturer’s instructions for amino acid analysis (Waters Corporation, Milford, MA, USA). Analysis was carried out as reported by Kruk et al. [29] with small modifications. Cell-free supernatants were deproteinated using 3 kDa cutoff filters (10,000 × g, 15 min), diluted with borate buffer, and derivatized using the AccQ-Tag reagents (Waters). The chromatographic separation with UHPLC system was performed as described previously [30]. Amino acids were quantified based on the internal standard method using L-norvaline as an internal standard. A six-point calibration curve for each amino acid was established by calculating the ratio of peak area to the area of norvaline.

#### Statistical analysis

Statistical analyses were performed using SPSS v24.0 (IBM). Differences between conditions were evaluated using unpaired t-tests with Bonferroni correction. Significance thresholds are indicated in figure legends.

### Label-free quantitative (LFQ) proteomics

#### Extraction of proteins

PFR cells were grown in five biological replicates on LAC, GLC, or FRC under the conditions described above and harvested at mid-log phase (LAC: 20 h, OD_600_ 0.54 ± 0.03; GLC: 24 h, OD_600_ 0.50 ± 0.01; FRC: 40 h, OD_600_ 0.66 ± 0.01). Cells were collected by centrifugation (5000 × g, 15 min, 4 °C), washed twice with ice-cold TEAB (pH 8.5) to remove loosely bound extracellular proteins [31], heated at 95 °C, and stored at -80 °C. For protein extraction, frozen cells were thawed at room temperature, suspended in 0.1% RapiGest SF (Waters Corp.) in 100 mM TEAB (pH 8.5), and transferred to FastPrep tubes containing ∼50 µL glass beads (ø 106 µm, Sigma Aldrich). Cells were disrupted in a FastPrep-24 (MP Biomedicals) at setting 6.5 for 4 × 30 s, centrifuged to pellet debris and beads, and frozen again at -80 °C. After thawing, samples were heated to 70 °C, cooled to room temperature, and subjected to a second round of FastPrep disruption. Then, 250 µL of 0.2% RapiGest in 100 mM TEAB (pH 8.0) was added, and samples were incubated at room temperature for 2-3 h with frequent mixing. Cell-free extracts were recovered by centrifugation (5000 × g, 5 min, RT), and protein concentration was measured using a NanoDrop 2000/2000c spectrophotometer (Thermo Fisher).

#### Proteomic analysis

Samples for LFQ proteomics were prepared using the PAC method [32] with modifications. Protein aggregation was induced from 10 µg of protein in RapiGest-buffer by adding acetonitrile to 70%, followed by capture on 10 µL MagReSyn Amine magnetic beads (LabLife Nordic AB, Finland). Bead-bound proteins were washed with 100% acetonitrile and 70% ethanol and briefly air-dried. For on-bead digestion, beads were submerged in 50 mM Tris-HCl (pH 8.0), proteins were reduced with 10 mM DTT for 30 min at 37 °C, alkylated with 20 mM iodoacetamide for 30 min at room temperature, and quenched with 20 mM DTT. Proteins were digested overnight at 37 °C with Trypsin Platinum (1:10, enzyme:protein) and rLysC (1:50)(Promega). After bead separation, peptides were acidified with 0.1% formic acid, purified using C18 microcolumns (Millipore), dried, resuspended in 0.1% formic acid, and analyzed by LC-MS/MS on a nanoElute UHPLC coupled to a timsTOF fleX mass spectrometer with a CaptiveSpray source (Bruker Daltonics, Bremen, Germany). Peptides were separated with a 60 min gradient at 300 nL/min, and data were acquired in DDA-PASEF mode using 10 MS/MS scans per cycle.

Raw MS files were processed in MaxQuant v2.0.1.0 [33,34] against the NZ_CP010341.1 database [25] containing 2,320 protein sequences. Carbamidomethylation of cysteine was set as a fixed modification, while methionine oxidation and protein N-terminal acetylation were included as variable modifications. Peptide mass tolerance was set to 20 ppm for the first search and 10 ppm for the main search. Trypsin without proline restriction was specified with up to two missed cleavages. Protein identification required at least one unique or razor peptide, and peptide and protein FDRs were controlled at 1% using reversed sequences. The raw proteomics data have been deposited to the ProteomeXchange Consortium via the PRIDE [35] partner repository with the dataset identifier PXD079229.

#### Proteome statistics

Downstream proteomic analyses were performed in Perseus v1.6.15.0 [36] using LFQ intensity values from the identified protein groups, with protein annotations confirmed by retrieving the protein sequences for the identified proteins from UniProtKB [37]. Decoys, contaminants, and proteins identified only by site were excluded. Proteins detected in at least three of five replicates in at least one condition were retained, data were log2 transformed and missing values were imputed from a normal distribution (width 0.3, downshift 1.8) and subjected to principal component analysis (PCA) to assess global proteomic variation and sample clustering between conditions. Z-score-normalized LFQ values were used for Euclidean hierarchical clustering and heatmap generation. Significant abundance changes were identified using multi-sample ANOVA with permutation-based FDR correction at 0.05, followed by Tukey’s HSD *post hoc* testing.

## RESULTS

### Growth dynamics and metabolite formation were strongly carbon source-dependent

**Figure 1** summarizes the experimental workflow designed to assess how carbon source shapes growth, metabolism, cofactor production, and proteome-wide adaptation in PFR DSM 20271. An initial screen across five carbon sources evaluated biomass formation, metabolic end-products, B12 production, and riboflavin uptake. Based on this screen, three representative conditions - lactate (LAC), glucose (GLC), and fructose (FRC) - were selected for LFQ proteomics, followed by bioinformatics and statistical analysis. These datasets then guided follow-up studies on B12 production and B2 uptake, organic acid production, amino acid uptake, and colony-forming ability. Growth kinetics differed markedly across substrates **(Figure 2A)**. Lactate supported the fastest growth rate (0.13 h⁻¹), reaching stationary phase after ∼50 h, whereas fructose showed the lowest rate (0.04 h⁻¹) and reached stationary phase after ∼90 h. Despite slower growth, GLC and FRC cultures reached the greatest final cell densities (OD₆₀₀ 6.1 ± 0.17 and 4.7 ± 0.26, respectively), compared with LAC (OD_600_ 2.4 ± 0.13) **(Table 1)**. Biomass recovery after 168 h followed the same trend. Substrate consumption and pH profiles reflected these growth patterns. LAC and GLC were largely depleted during the anaerobic phase, whereas FRC was consumed more slowly. Higher biomass formation in hexose cultures coincided with medium acidification (pH 4.7-5.0), whereas LAC cultures maintained near-neutral pH (∼6.4). During anaerobic growth, LAC and GLC cultures produced the highest levels of propionate and acetate, whereas FRC cultures produced substantially lower concentrations. Transition to microaerophilic conditions shifted fermentation toward higher acetate-to-propionate ratios in all cultures.

**Figure 1.**
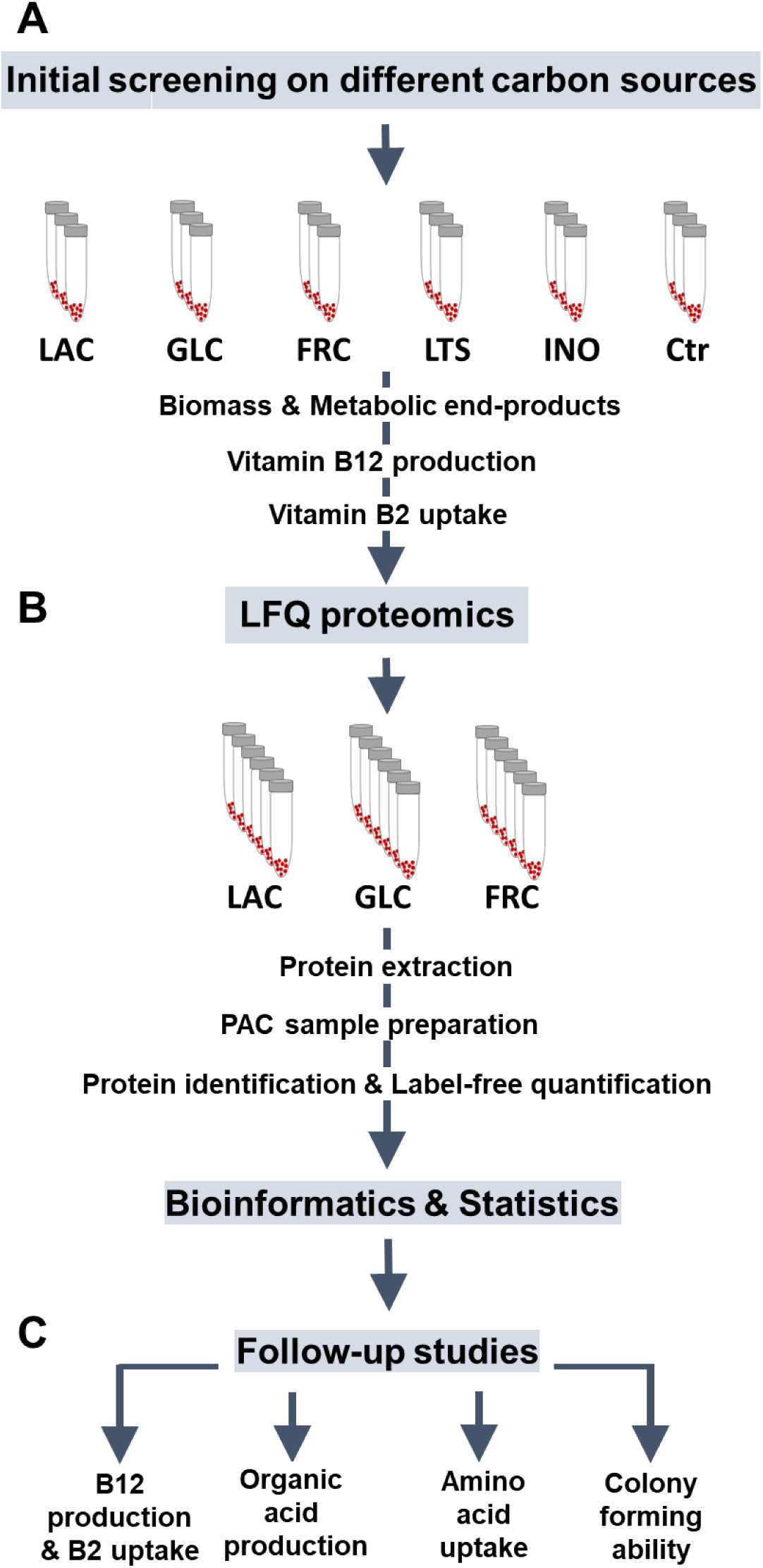
Experimental workflow for evaluating carbon source-dependent effects on growth, metabolism, cofactor production, and proteome-wide adaptation in PFR DSM 20271. **(A)** Initial screening across five carbon sources - lactate (LAC), glucose (GLC), fructose (FRC), lactose (LTS), and inositol (INO) - was used to assess biomass formation, metabolic end-products, vitamin B12 production, and riboflavin (vitamin B2) uptake. **(B)** Based on the screening results, LAC-, GLC-, and FRC-grown cultures were selected for label-free quantitative (LFQ) proteomics. Cells were processed by protein extraction, PAC-based sample preparation, protein identification, and label-free quantification, followed by bioinformatics and statistical analysis. (**C)** Proteomic and phenotypic findings then guided follow-up studies of B12 production and B2 uptake, organic acid production, amino acid uptake, and colony-forming ability. Ctr indicates control medium without added carbon substrate.

**Figure 2.**
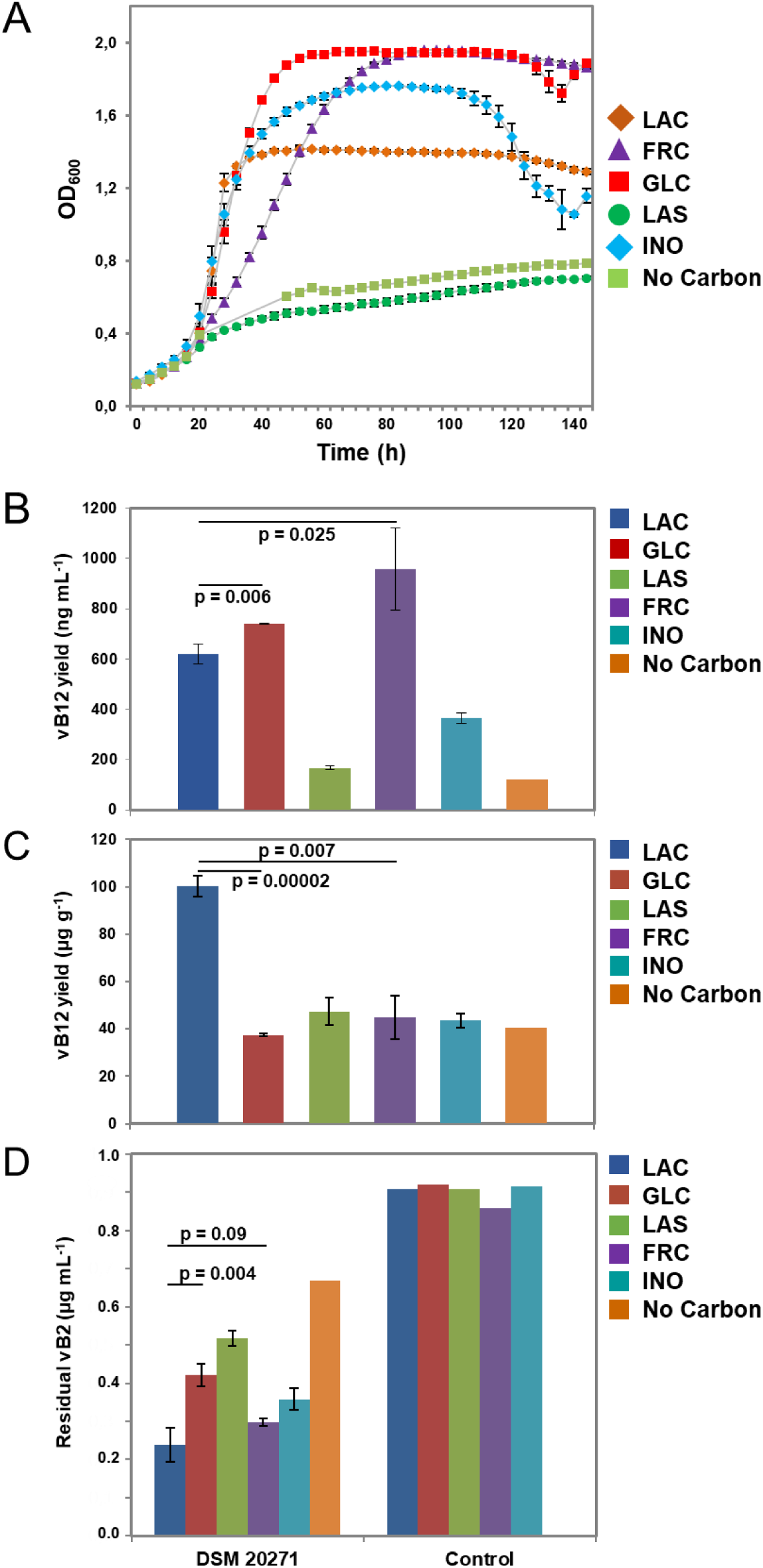
Growth characteristics of PFR and related B12 production. **(A)** Growth of the PFR strain DSM 20271 in yeast extract-tryptone (YE)-based medium supplemented with different carbon sources and using a Bioscreen to monitor cell density at 600 nm at the indicated timepoints. **(B)** Total B12 production yields (µg mL^−1^ of total culture volume) reached by PFR in YE containing different carbon sources. **(C)** B12 synthesis efficiency (µg g^−1^ of wet cell biomass) of PFR grown in YE with different carbon sources. Error bars indicate the standard deviation of three independent replicate analyses. Unpaired t-test with equal variance assumption indicates statistical significance between the indicated samples.

**Table 1.** Growth characteristics, production of metabolites (propionate and acetate), residual carbon in spent culture supernatants, and wet cell mass of *P. freudenreichii strain* DSM20271^T^ grown in yeast extract-tryptone-based medium with different carbon sources.

| Parameters | TIME (h) | CARBON SOURCE |  |  |  |  |  |
| --- | --- | --- | --- | --- | --- | --- | --- |
|  |  | Lactate | Glucose | Lactose | Fructose | Inositol | No carbon |
| Optical density (OD<sub>600</sub>) | 72 | 1.8 (0.01) | 6.3 (0.18) | 0.5 (0.02) | 3.3 (0.06) | 3.7 (0.01) | 0.5 |
|  | 168 | 2.4 (0.13) | 6.1 (0.17) | 1.5 (0.02) | 4.7 (0.26) | 3.7 (0.14) | 1.5 |
| pH | 72<sup>a</sup> | >6.1 | 4.7 | >6.1 | 5.6 | 4.7 | >6.1 |
|  | 168<sup>b</sup> | 6.4 (0.05) | 4.7 (0.01) | 6.5 (0.03) | 5.0 (0.01) | 4.8 (0.01) | 6.5 |
| Propionate (mg/mL) | 72 | 3.4 (0.06) | 3.2 (0.08) | 0.3 (0.01) | 0.9 (0.07) | 1.9 (0.05) |  |
|  | 168 | 2.3 (0.11) | 2.3 (0.06) | 0.2 (0.00) | 1.4 (0.17) | 1.8 (0.13) | 0.2 |
| Acetate (mg/mL) | 72 | 1.4 (0.04) | 1.2 (0.02) | 0.1 (0.01) | 0.5 (0.03) | 1.0 (0.03) |  |
|  | 168 | 1.7 (0.11) | 1.3 (0.03) | 0.2 (0.02) | 1.3 (0.10) | 1.4 (0.12) | 0.3 |
| Residual carbon (mg/mL) | 72 | <0.10 | 0.10 | 8.3 (0.1) | 5.0 (0.1) | 1.3 (0.0) |  |
|  | 168 | <0.10 | <0.10 | 7.4 (0.4) | 1.1 (0.1) | 0.0 (0.0) |  |
| Wet cell mass (g/20 mL) | 168 | 0.11 (0.00) | 0.36 (0.04) | 0.06 (0.01) | 0.40 (0.03) | 0.16 (0.00) | 0.06 |

### Carbon source determines volumetric and specific B12 production

Volumetric B12 production varied strongly by carbon source **(Figure 2B)**. Without carbon supplementation, DSM 20271 produced 120 ng mL^−1^ B12. LAC and GLC increased yields to 619 and 741 ng mL^−1^, respectively, whereas FRC supported the highest volumetric production (957 ng mL^−1^). In contrast, biomass-normalized values showed the opposite trend **(Figure 2C)**. LAC-grown cells exhibited the highest specific B12 content (∼100 µg g^−1^ fresh biomass), whereas values for the other substrates remained below 50 µg g^−1^.

Because flavin cofactors derived from riboflavin participate in redox reactions associated with corrinoid biosynthesis and central carbon metabolism, residual riboflavin was monitored as an indicator of cofactor demand during B12 production. Residual riboflavin decreased during fermentation **(Figure 2D)**. LAC cultures showed the lowest residual levels (0.24 µg mL^−1^), indicating high net uptake from the medium, whereas FRC cultures retained higher levels (0.56 µg mL^−1^) despite higher volumetric B12 production. Together, these findings show that carbon source influences both total B12 production and biomass-specific corrinoid accumulation, while also altering riboflavin utilization.

### Proteomic adaptation to lactate and hexose growth conditions

Label-free quantitative proteomic analysis identified 1,560 proteins across PFR cells grown under LAC, GLC, and FRC conditions **(Table S1)**, with high reproducibility across biological replicates **(Figure S1)**. Multivariate analyses, including principal component analysis **(Figure 3A)** and hierarchical clustering **(Figure 3B)**, showed clear group separation according to carbon source. Proteome-wide ANOVA following Perseus processing identified 851 proteins with significantly different abundances among growth conditions (q < 0.05) (**Table S2**). The greatest abundance differences are summarized in **Table 2**.

**Figure 3.**
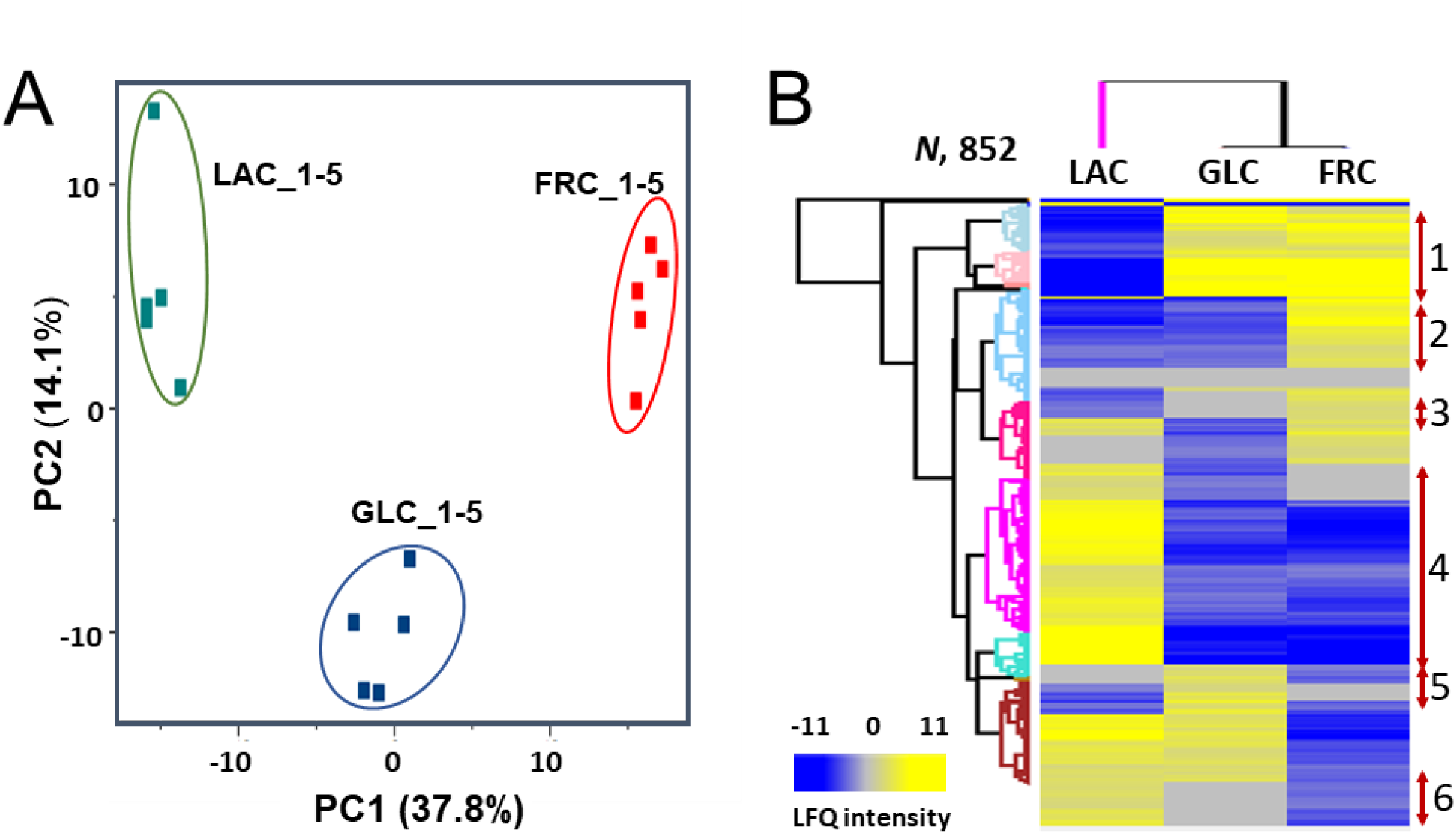
Multivariate analyses of LFQ identification data. **(A)** Principal component analysis (PCA) of all detected proteins based on LFQ intensities. The five biological replicates per condition cluster closely, indicating high reproducibility and clear separation of proteomes according to carbon source (LAC, GLC, FRC). **(B)** Hierarchical clustering of proteins with significant abundance differences across conditions. Proteome-wide ANOVA (permutation-based FDR = 0.05; q < 0.05), followed by post hoc Tukey’s HSD test, identified 851 differentially abundant proteins. Z-score-normalized LFQ intensities were subjected to Euclidean distance-based clustering, resulting in six distinct clusters that capture carbon source-dependent patterns and relationships within the LAC-, GLC-, and FRC-associated proteomes.

**Table 2.**
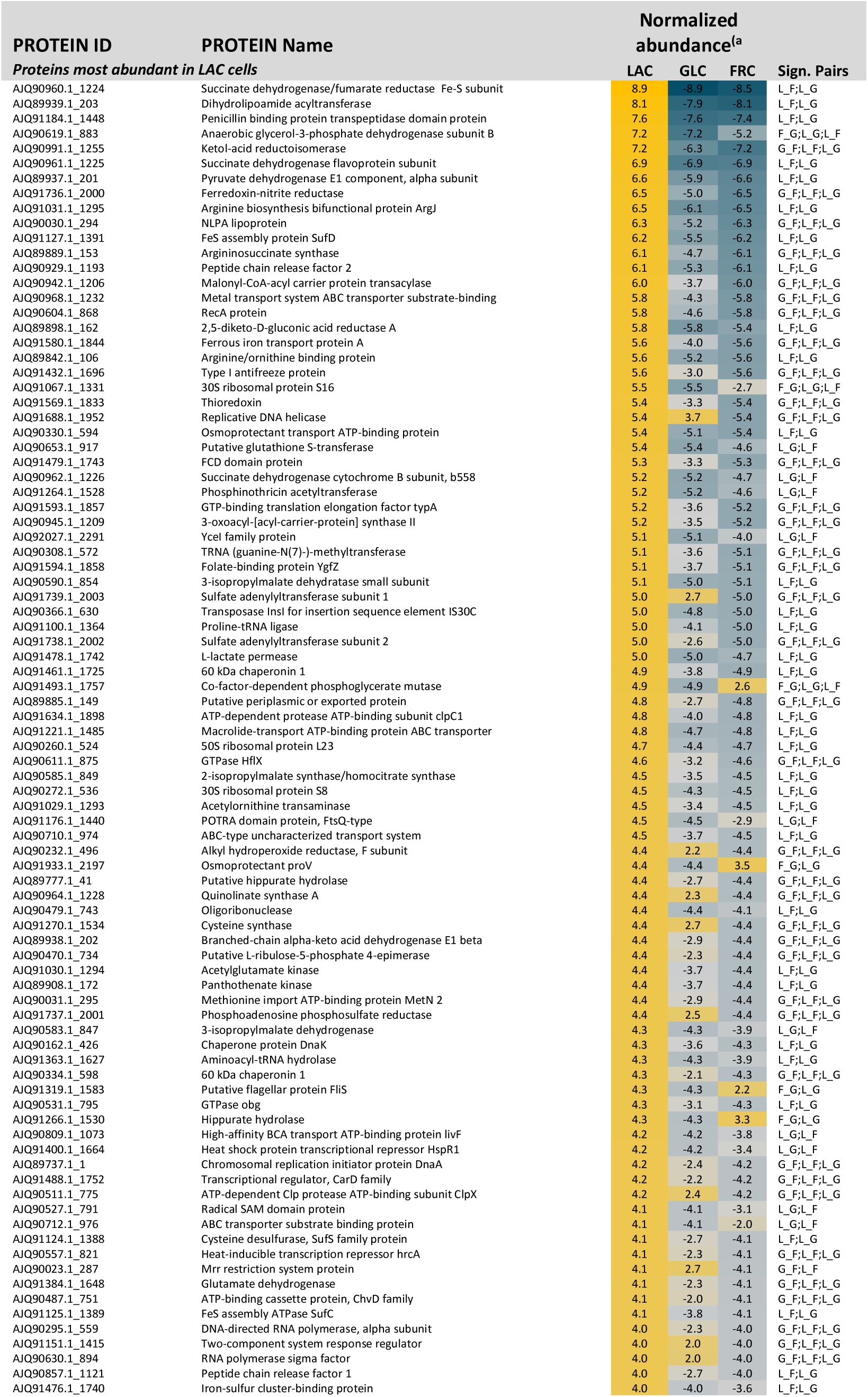
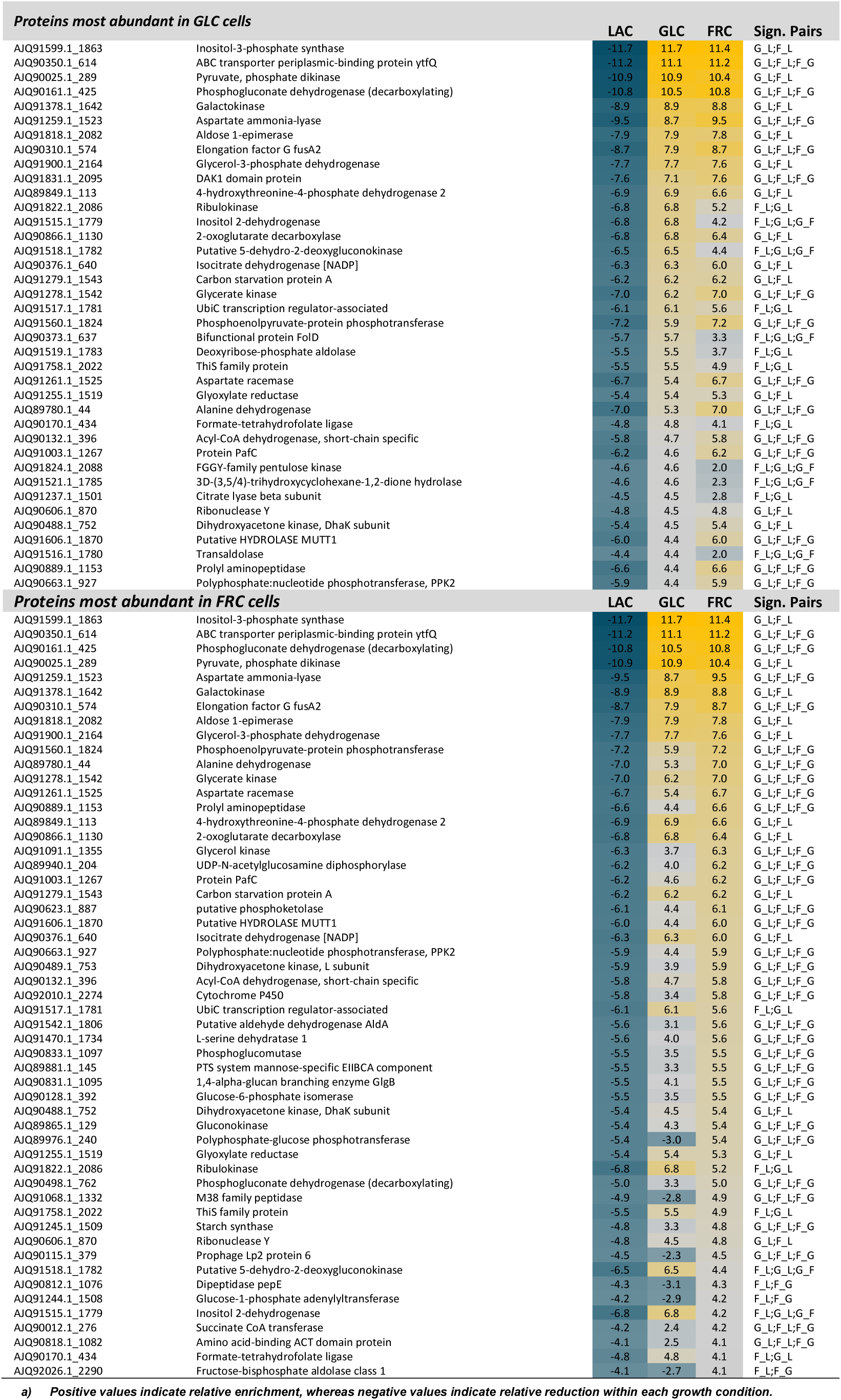
Selected proteins with significant abundance change in LAC (L), GLC (G) and FRC (F) cells.

#### LAC-associated changes

In LAC-grown cells, proteins with increased abundance were mainly associated with respiratory metabolism, redox homeostasis, amino acid biosynthesis, transport, and stress adaptation. Enrichment of respiratory electron-transfer components, including SdhB/FrdB (normalized abundance = 8.9) and SdhA/FrdA (6.9), and anaerobic glycerol-3-phosphate dehydrogenase subunit GlpB (7.2), together with pyruvate oxidation enzymes such as AceF/DlaT (8.1) and AceE/PdhE1 (6.6), indicated enhanced respiratory activity during growth on lactate. Increased abundance of thioredoxin (5.4), ferredoxin-nitrite reductase NirA (6.5), and Fe-S cluster assembly proteins SufD, SufS, and SufC further supported elevated redox and cofactor-related activity. LAC-grown cells also showed increased levels of the lactate permease LldP, metal transport proteins including FeoA, and osmoprotectant transport systems, suggesting coordinated adaptation to lactate utilization and cofactor demand. In addition, higher abundance of molecular chaperones and replication-associated proteins, including GroEL, DnaK, RecA, and DnaA, pointed to increased biosynthetic activity and stress-responsive capacity in LAC-grown cells.

#### FRC-associated changes

FRC-grown cells showed a distinct enrichment of proteins involved in fructose and glucose interconversion, sugar phosphate turnover, pentose metabolism, and carbohydrate storage. Increased abundance of phosphoglucomutase, glucose-6-phosphate isomerase, glucose-1-phosphate adenylyltransferase, fructose-bisphosphate aldolase, ribulokinase, and gluconokinase indicated enhanced metabolic flexibility during fructose utilization. In addition, enrichment of starch synthase and GlgB suggested a fructose-specific shift toward carbohydrate reserve metabolism. Proteins involved in phosphate and energy metabolism, including PPK2, polyphosphate-glucose phosphotransferase, and phosphoenolpyruvate-protein phosphotransferase, were also increased, consistent with elevated energetic demand during rapid sugar phosphate turnover and carbohydrate catabolism.

#### Hexose associated changes

GLC- and FRC-grown cells showed strong enrichment of proteins associated with carbohydrate uptake, central carbon metabolism, energy generation, and reductive metabolism. High abundance of the ABC transporter component YtfQ (11.1 in GLC; 11.2 in FRC), together with Ino1 (11.7; 11.4), PpdK (10.9; 10.4), and Gnd (10.5; 10.8), indicated increased carbohydrate uptake, glycolytic activity, and pentose phosphate pathway flux under both hexose conditions. Additional enrichment of GalK, GalM, GlpD, transaldolase, and phosphoenolpyruvate-protein phosphotransferase further supported enhanced carbohydrate interconversion and redox-associated metabolism during growth on glucose and fructose. Hexose-grown cells also showed increased abundance of proteins linked to overflow metabolism, alternative carbon utilization, and reductive metabolism, particularly under FRC conditions. Enrichment of AspA, alanine dehydrogenase, glycerate kinase, phosphoketolase-related proteins, and glyoxylate reductase suggested extensive redistribution of carbon and nitrogen metabolism under hexose conditions. Consistently, GLC- and FRC-grown cultures produced approximately two-fold higher succinate levels than LAC-grown cultures (p < 2.0 × 10⁻⁶) **(Figure 4A)**, supporting increased reductive TCA-associated flux during hexose metabolism.

**Figure 4.**
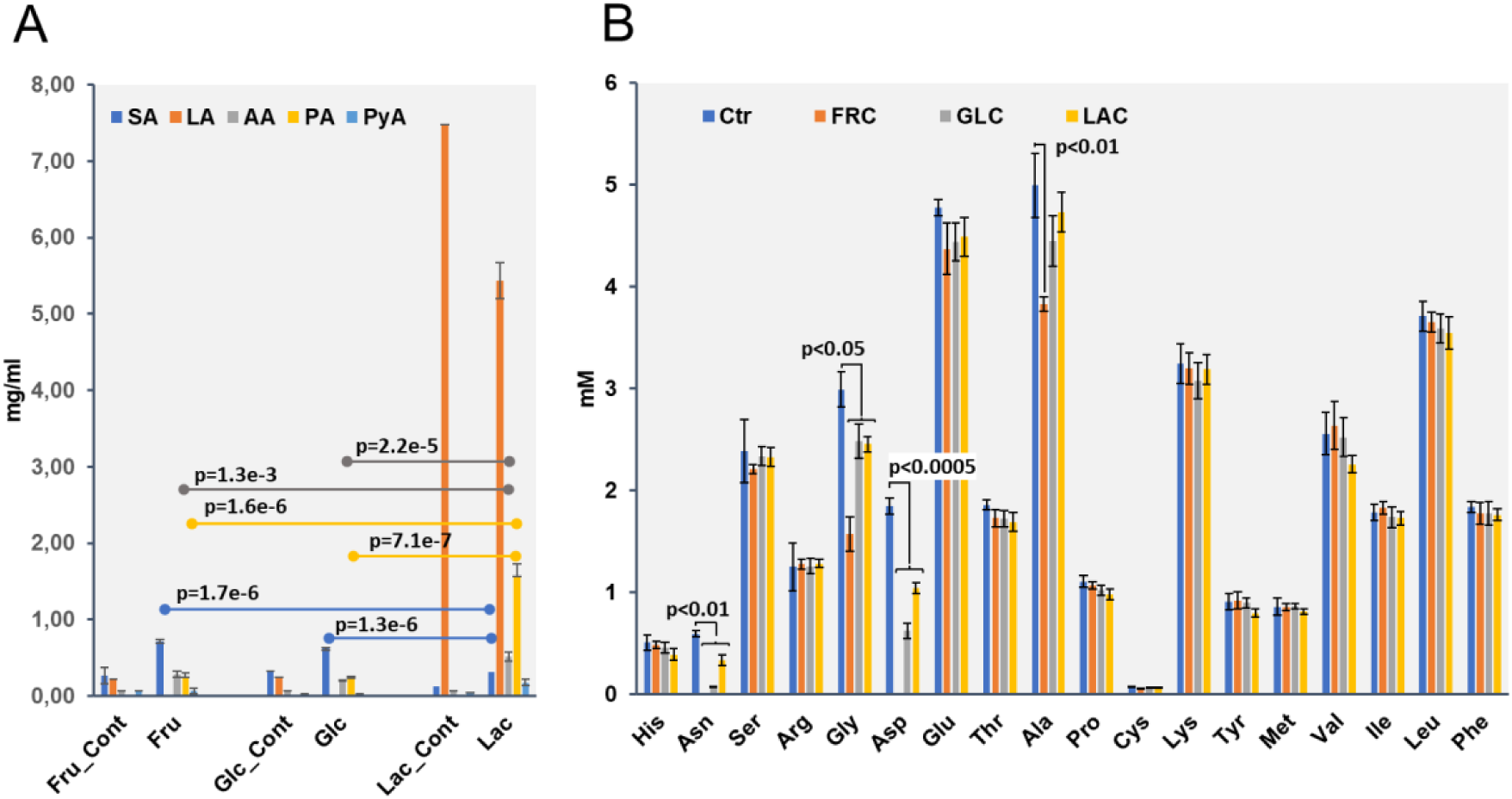
Organic acid production and amino acid consumption depend on carbon source. **(A)** Organic acid production by PFR grown on fructose (FRC), glucose (GLC), or lactate (LAC), compared to control medium (Ctr). Concentrations of succinic acid (SA), lactic acid (LA), acetic acid (AA), propionic acid (PA), and pyruvic acid (PyA) were determined in cell-free supernatants by HPLC after cultivation. **(B)** Amino acid levels in spent culture supernatants following growth on FRC, GLC, or LAC. Amino acids were quantified after derivatization using UHPLC. Hexose-grown cultures showed marked depletion of several amino acids compared with LAC-grown cultures, consistent with increased amino acid utilization under these conditions. Statistical significance was determined using unpaired t-tests with Bonferroni correction.

Together, these findings indicate that lactate promotes a respiratory and stress-adapted proteomic state, whereas glucose and fructose favor rapid carbohydrate turnover, overflow metabolism, and extensive metabolic reorganization.

### Carbon source redirects amino acid and nitrogen metabolism

Proteomic analysis also revealed marked carbon source-dependent differences in amino acid and nitrogen metabolism **(Table 2)**. LAC-grown cells showed increased abundance of proteins associated with amino acid biosynthesis, sulfur assimilation, and Fe-S cluster assembly. Among the most enriched proteins were arginine biosynthesis bifunctional protein ArgJ (normalized abundance = 6.5), ferredoxin-nitrite reductase NirA (6.5), ketol-acid reductoisomerase IlvC (7.2), argininosuccinate synthase (6.1), and acetylglutamate kinase (4.4), indicating enhanced branched-chain and arginine biosynthetic activity during growth on lactate. Increased abundance of sulfur-associated proteins, including sulfate adenylyltransferase subunits 1 and 2, phosphoadenosine phosphosulfate reductase, and cysteine synthase, together with Fe-S assembly proteins (SufD, SufS, SufC), further supported enhanced sulfur utilization and cofactor biosynthesis under lactate conditions.

In contrast, GLC- and FRC-grown cells displayed increased abundance of proteins associated with amino acid catabolism, carbon redistribution, and reductive metabolism. Aspartate ammonia-lyase AspA showed strong enrichment under GLC (8.7) and FRC (9.5) conditions relative to LAC-grown cells. Additional enrichment of alanine dehydrogenase, aspartate racemase, glycerate kinase, and glyoxylate reductase suggested coordinated redistribution of carbon and nitrogen metabolism during growth on hexoses. Hexose-grown cells also showed elevated levels of phosphogluconate dehydrogenase Gnd, pyruvate phosphate dikinase PpdK, and glycerol-3-phosphate dehydrogenase GlpD, consistent with elevated pentose phosphate pathway activity, increased NADPH demand, and enhanced reductive metabolism during growth on glucose and fructose.

### Hexose-associated amino acid depletion is linked to reduced aerobic viability

To validate the proteomic findings, amino acid profiles were analyzed during growth on different carbon sources. Asparagine, aspartate, and glycine were rapidly depleted in hexose-grown cultures **(Figure 4B)**, supporting increased amino acid utilization under these conditions. Although colony-forming ability in liquid cultures remained comparable among treatments, colony formation on solid media was strongly carbon source-dependent. GLC- and FRC-containing plates showed markedly reduced colony formation, particularly under semiaerobic and aerobic conditions **(Figure 5A)**. Notably, aspartate supplementation restored colony formation on FRC plates **(Figure 5B)**, linking hexose-associated amino acid depletion to impaired aerobic viability. Thus, hexose metabolism was associated with increased amino acid consumption, particularly of aspartate and asparagine, together with reduced colony-forming ability under oxygen-exposed conditions.

**Figure 5.**
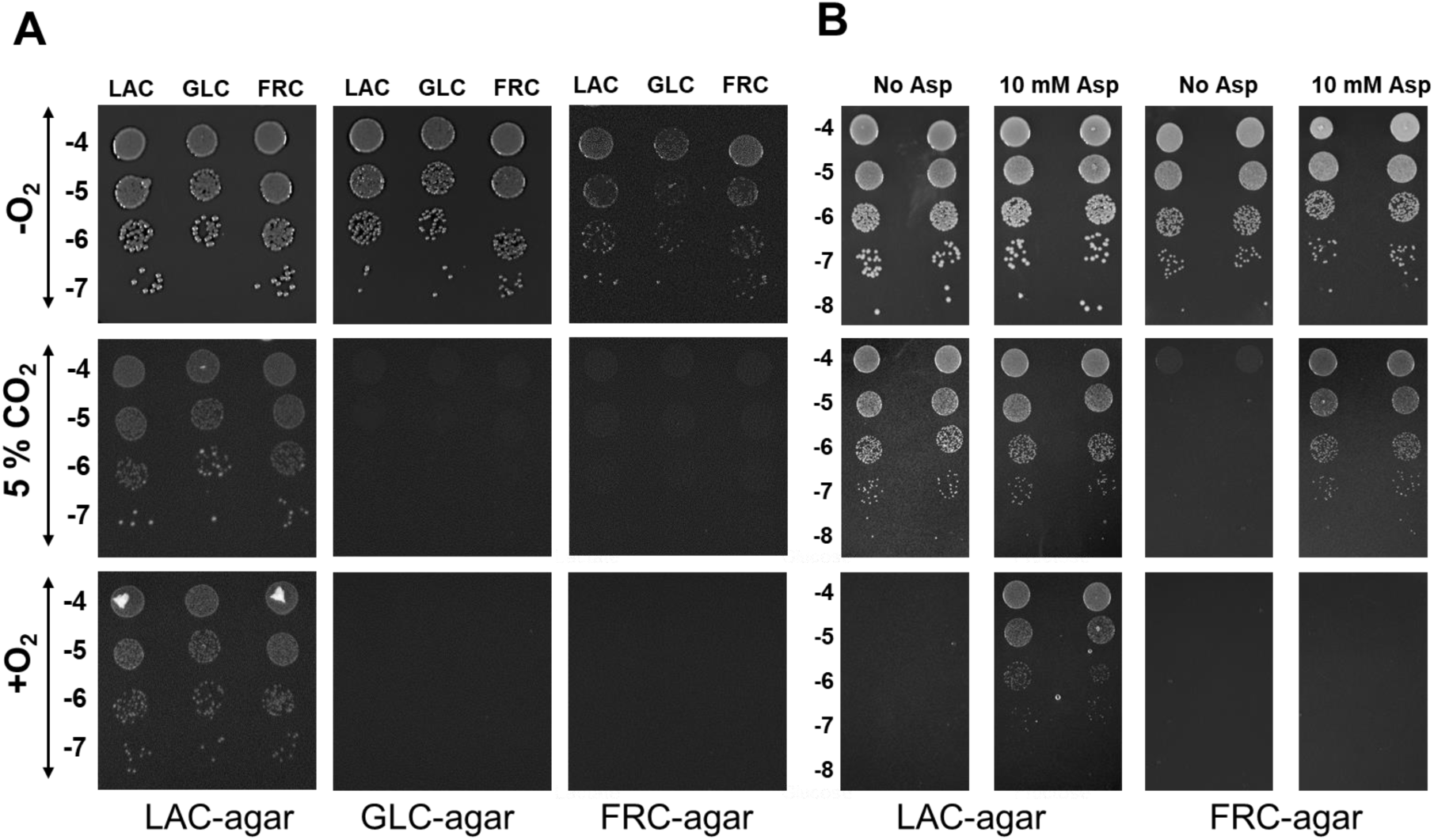
Carbon source- and oxygen-dependent colony-forming ability of PFR. **(A**) Colony formation of cells grown on lactate (LAC), glucose (GLC), or fructose (FRC) and spotted as serial dilutions (10⁻⁴ to 10⁻⁷/10⁻⁸) on YE agar containing the indicated carbon source. Plates were incubated at 30 °C under anaerobic (-O₂; 4 days), semiaerobic (5% CO₂; 6 days), or aerobic (+O₂; 8 days) conditions. Colony formation was comparable across preculture conditions when plated on LAC agar, whereas growth on GLC and FRC agar resulted in reduced colony formation, particularly under oxygen-exposed conditions. **(B)** Effect of aspartate supplementation on colony formation. Cells grown on LAC or FRC were spotted on YE agar containing the indicated carbon source with or without aspartate (10 mM). Anaerobic and semiaerobic plates were incubated as in panel A, whereas aerobic plates were incubated for 6 days. Aspartate supplementation restored colony formation on FRC agar, particularly under semiaerobic conditions, whereas only minor effects were observed on LAC agar. These results support a role for aspartate in maintaining growth and viability under redox-constrained conditions associated with fructose metabolism.

### Aspartate metabolism drives succinate accumulation as an alternative electron sink

Because proteomic analysis revealed strong enrichment of AspA together with increased succinate production under hexose conditions, we investigated whether aspartate contributes to reductive metabolism through fumarate-dependent succinate formation. To test this, cultures were supplemented with 10 mM aspartate, which triggered a 3-4-fold increase in succinate accumulation during anaerobic growth in both FRC+ASP and LAC+ASP cultures (0.29 and 0.34 mg mL^−1^, respectively) compared with unsupplemented cultures (∼0.08-0.10 mg mL^−1^) **(Figure 6A)**. This is consistent with rapid aspartate depletion and strong upregulation of aspartate ammonia-lyase, indicating active conversion of aspartate to fumarate and subsequent reduction to succinate. Following the shift to microaerophilic conditions, succinate was rapidly consumed in FRC+ASP cultures but remained stable in LAC+ASP cultures. Aspartate supplementation did not significantly affect propionate or acetate production. Together, these findings support the interpretation that aspartate primarily supports maintenance of redox homeostasis rather than as a major carbon source, supporting aerobic viability under hexose metabolism.

**Figure 6.**
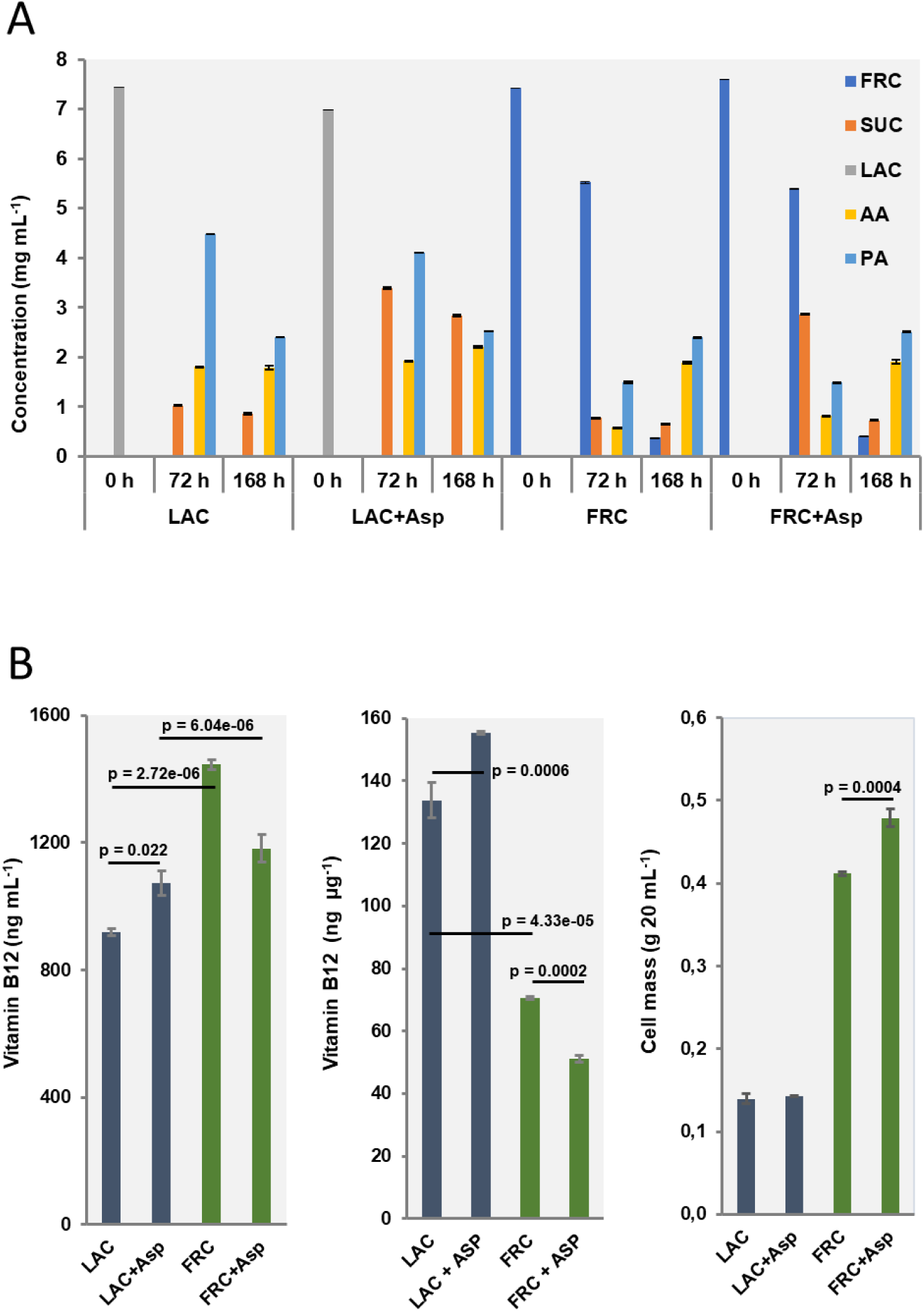
Aspartate supplementation reshapes redox metabolism and vitamin B12 production. **(A)** Time-resolved metabolite profiles in PFR cultures grown on lactate (LAC) or fructose (FRC), with or without aspartate supplementation (10 mM). Concentrations of fructose (FRC), lactate (LAC), succinate (SUC), acetate (AA), and propionate (PA) were measured at 0, 72, and 168 h. Aspartate supplementation led to increased succinate accumulation during anaerobic growth, particularly in FRC cultures, consistent with activation of an aspartate-linked fumarate–succinate pathway. Residual substrate consumption and organic acid profiles differed between LAC and FRC conditions. **(B)** Effects of aspartate supplementation on volumetric B12 production (left), biomass-normalized (specific) B12 production (middle), and biomass (right) in LAC- and FRC-grown cultures. Aspartate supplementation increased volumetric and specific B12 production in LAC cultures but reduced both parameters in FRC cultures, while increasing cell biomass under FRC conditions. Statistical significance was determined using unpaired t-tests with Bonferroni correction.

### Aspartate supplementation exerts opposing effects on B12 production

Because B12 biosynthesis depends on both precursor availability and cellular redox balance, we next examined whether aspartate supplementation differentially affects B12 production during lactate and fructose metabolism. Aspartate supplementation increased B12 production in LAC-grown cells, improving both volumetric and specific yields by 16-17%, but reduced production in FRC-grown cultures by 18% volumetrically and 27% on a biomass-specific basis **(Figure 6B)**. These opposing responses suggest that aspartate supports B12 precursor supply under lactate metabolism, whereas during fructose metabolism it is preferentially diverted toward maintenance of cellular redox homeostasis. Aspartate availability modulates B12 biosynthesis in a carbon source-dependent manner, enhancing production during lactate metabolism but limiting it during fructose-driven redox balancing.

### Carbon source differentially affects corrinoid- and riboflavin-associated pathways

Proteomic analysis revealed distinct carbon source-dependent abundance patterns in proteins associated with vitamin B12 biosynthesis, cobalt transport, and riboflavin metabolism **(Figure 7; Table S2)**. Several enzymes involved in cobalamin biosynthesis showed differential enrichment across conditions, indicating carbon-specific regulation of B12 assembly pathways. Early-stage B12 biosynthesis enzymes, including CobA, CobA2, and the intermediate pathway enzyme CbiC, were most abundant under FRC conditions, consistent with the high volumetric B12 production observed during fructose metabolism. In contrast, the late-stage biosynthetic enzyme CobQ was most enriched in GLC-grown cells, whereas HemD and CbiX showed higher abundance in both GLC- and LAC-grown cultures relative to FRC, suggesting differential regulation of tetrapyrrole and cobalt-associated biosynthetic steps depending on the carbon source.

**Figure 7.**
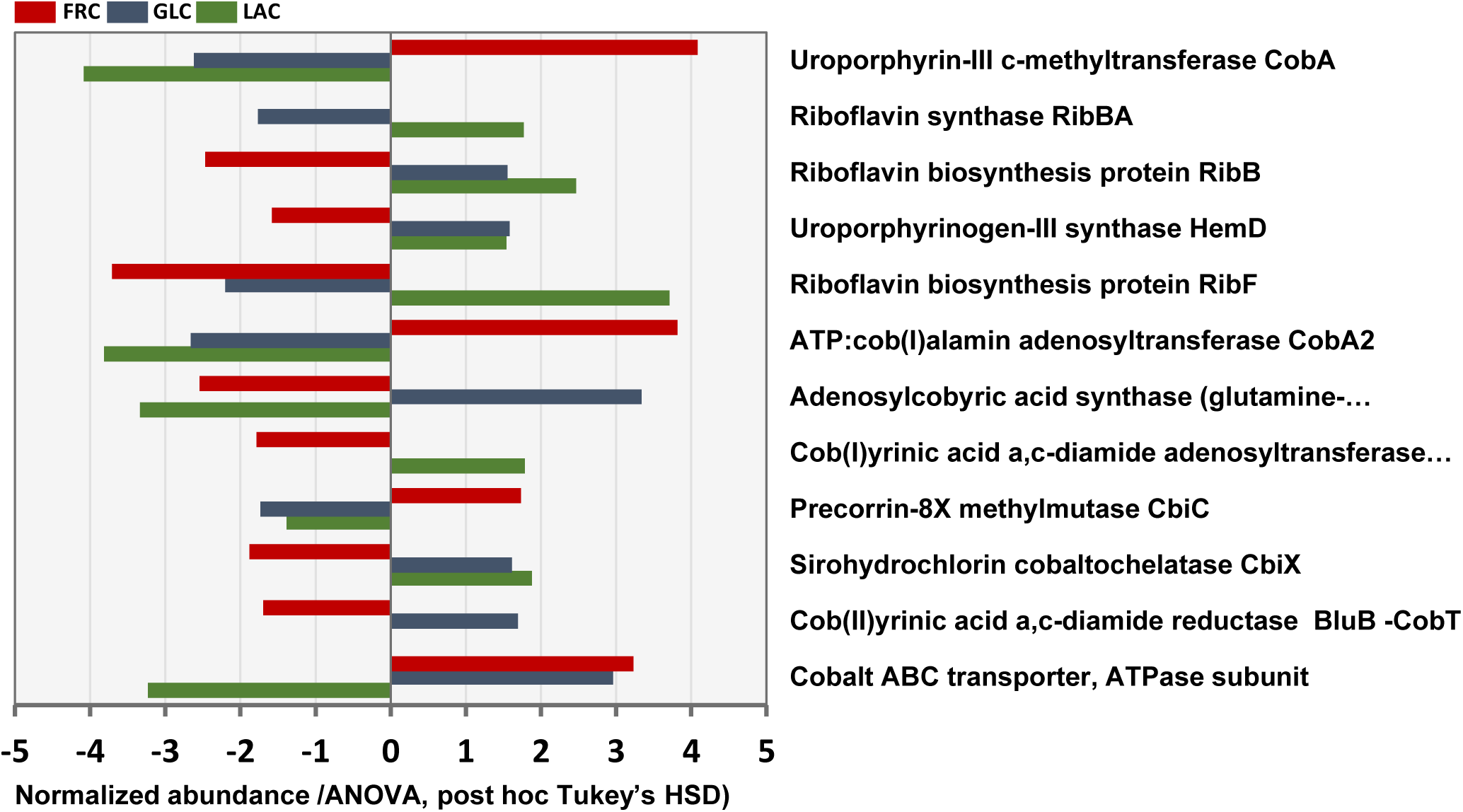
Carbon source-dependent abundance patterns of vitamin B12 biosynthesis, cobalt transport, and riboflavin pathway proteins. Proteins associated with *de novo* B12 biosynthesis, cobalt transport, and riboflavin metabolism showing significant abundance differences across lactate (LAC), glucose (GLC), and fructose (FRC) conditions. Values represent log₂-transformed abundance differences derived from proteome-wide ANOVA analysis. Early corrin ring assembly and adenosylation enzymes (e.g., CobA, CobA2, CobO, CbiC) and the mid-pathway enzyme precorrin-8X methylmutase (CbiC) show carbon source-dependent variation, with several enriched under hexose conditions. In contrast, the late-stage amidation enzyme CobQ and early tetrapyrrole pathway enzymes (e.g., HemD, CbiX) display distinct patterns across substrates. Cobalt ABC transporter components are more abundant in hexose-grown cells, whereas riboflavin biosynthesis enzymes (RibBA, RibB, RibF) show higher abundance under lactate conditions. The terminal lower ligand synthesis enzyme (BluB-CobT) exhibits independent variation, indicating partial decoupling between precursor biosynthesis and final B12 assembly.

Proteins associated with cobalt uptake were similarly enriched under hexose conditions. Cobalt ABC transporter components displayed increased abundance in both GLC- and FRC-grown cells, supporting elevated trace metal demand during enhanced corrinoid biosynthesis under carbohydrate metabolism. In contrast, proteins involved in riboflavin biosynthesis and flavin metabolism were preferentially enriched during growth on lactate. RibBA showed the strongest response, with ∼2.7-fold higher abundance in LAC relative to FRC cultures, whereas RibF and the riboflavin synthase alpha subunit RibB were elevated by ∼1.7-fold and ∼1.6-fold, respectively. These patterns are consistent with increased flavin cofactor generation and redox-associated metabolic activity during lactate metabolism, in agreement with the low residual riboflavin levels measured in LAC cultures.

Notably, the terminal B12 biosynthesis enzyme BluB-CobT2 displayed condition-specific regulation distinct from upstream pathway components, suggesting partial decoupling between precursor synthesis and final cobalamin assembly.

### Model of carbon source-dependent balancing between redox homeostasis and B12 biosynthesis in PFR

The concluding model **(Figure 8)** was developed by integrating physiological, metabolite, amino acid, and LFQ proteomic datasets from LAC-, GLC-, and FRC-grown cells. Growth, metabolite, amino acid, vitamin, viability, and proteomic datasets were integrated to define carbon source-dependent physiological configurations. Collectively our findings support a model in which PFR allocates metabolic resources between redox balancing and B12 biosynthesis according to carbon source. LAC promotes a respiratory, redox-balanced state associated with amino acid biosynthesis, flavin metabolism, and high biomass-specific B12 accumulation. In contrast, GLC and FRC induce a redox-constrained state characterized by increased carbohydrate turnover, pentose phosphate pathway activity, and succinate formation through the aspartate-to-fumarate pathway, which serves as an alternative electron sink. Under hexose conditions, aspartate is preferentially diverted toward redox balancing rather than corrinoid precursor supply, explaining its contrasting effects on B12 production in LAC- and FRC-grown cells. Thus, the prevailing physiological configuration determines the balance between redox homeostasis and efficient corrinoid biosynthesis.

**Figure 8.**
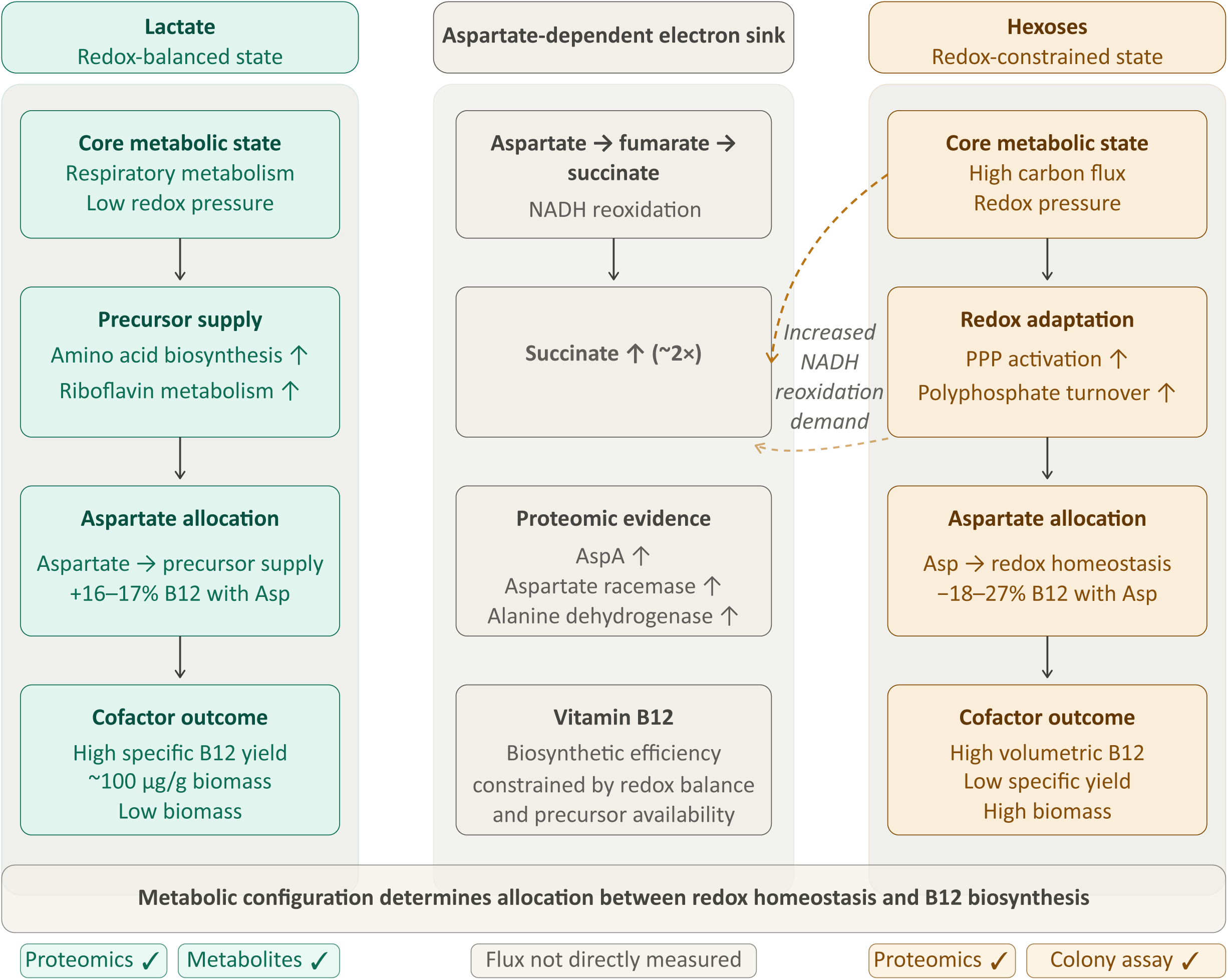
Model of carbon source-dependent metabolic configurations linking redox homeostasis and vitamin B12 biosynthesis in PFR. Lactate (LAC) metabolism promotes a respiratory, comparatively redox-balanced configuration associated with efficient precursor supply, flavin metabolism, and high biomass-specific vitamin B12 production. In contrast, glucose and fructose (GLC/FRC) establish a high-flux, redox-constrained configuration characterized by increased carbohydrate turnover, pentose phosphate pathway (PPP) activity, and redistribution of carbon and reducing power toward maintenance of redox homeostasis. A proposed feature of this configuration is activation of an aspartate-dependent electron sink, in which aspartate is converted to fumarate and subsequently reduced to succinate. Under specific conditions, fumarate may function as a terminal electron acceptor, redirecting carbon flux from propionate formation toward acetate and CO₂ production. Together, the model illustrates how carbon source influences flux distribution, resource allocation, and redox homeostasis, with vitamin B12 biosynthesis emerging as a system-level property of the prevailing metabolic configuration rather than biosynthetic enzyme abundance alone.

## DISCUSSION

### Carbon source defines distinct physiological configurations

Here, we show that carbon source is a major determinant of metabolic organization in PFR, coordinating central carbon metabolism, redox homeostasis, amino acid utilization, and cofactor biosynthesis into distinct physiological states. Integration of physiological, metabolite, amino acid, and proteomic datasets revealed that lactate and hexoses establish fundamentally different metabolic configurations that differentially allocate resources between respiratory metabolism, redox balancing, and vitamin B12 biosynthesis (Figure 8). These observations are consistent with genome-scale modelling studies indicating that metabolic output in PFR is limited by the interplay between carbon flux distribution, energy generation, and redox balance, which together define feasible physiological states and pathway utilization [6]. Combined with previous transcriptomic analyses showing extensive metabolic reprogramming during nutrient limitation and stationary-phase adaptation in *P. freudenreichii* [21,38], our findings support a multi-layered regulatory framework in which carbon source and oxygen availability impose interconnected constraints on metabolism. Oxygen availability primarily influenced cofactor biosynthesis and stress adaptation, whereas carbon source defined the prevailing metabolic mode governing flux redistribution, redox handling, and biosynthetic prioritization.

### Hexose metabolism creates a redox-constrained, high-flux state

Hexose metabolism established a redox-constrained, high-flux metabolic state characterized by increased carbohydrate turnover, pentose phosphate pathway activity, overflow metabolism, and activation of alternative redox-balancing routes. Proteomic enrichment of PPP-associated enzymes, glycolytic proteins, phosphoketolase-related pathways, and polyphosphate-utilizing systems indicates elevated demand for reducing power generation and rapid carbon redistribution under glucose and fructose metabolism [39,40]. Increased abundance of polyphosphate kinase 2 (PPK2) and polyphosphate-glucose phosphotransferase further suggests enhanced turnover of intracellular polyphosphate. In *P. freudenreichii*, polyphosphate functions as a phosphate and energy storage polymer whose accumulation and utilization are strongly influenced by carbon source. Lactate-grown cells accumulate large amounts of long-chain polyphosphate in volutin granules, whereas glucose-grown cells contain substantially lower levels, likely due to continuous utilization during carbohydrate phosphorylation [41]. Although polyphosphate was not measured directly in the present study, the increased abundance of polyphosphate-associated proteins is consistent with enhanced phosphate and energy cycling accompanying rapid sugar-phosphate turnover under hexose metabolism.

In parallel, several observations point to increased reliance on an aspartate-dependent route for redox balancing. Hexose-grown cultures exhibited increased succinate production, strong induction of aspartate ammonia-lyase, rapid depletion of extracellular aspartate, and reduced aerobic colony-forming ability. Together, these findings support a model in which aspartate is converted to fumarate and subsequently reduced to succinate, thereby providing an alternative electron sink during periods of high carbon flux. This interpretation is consistent with biochemical evidence linking aspartate deamination to fumarate reduction and succinate formation [42]. Importantly, restoring aerobic viability via aspartate supplementation provides experimental evidence supporting a role for aspartate-dependent redox balancing in survival during hexose metabolism.

### Lactate supports respiratory metabolism and efficient cellular B12 accumulation

In contrast, lactate promoted a respiratory and comparatively redox-balanced physiological state associated with amino acid biosynthesis, sulfur assimilation, Fe-S cluster assembly, and flavin-associated metabolism. Enrichment of respiratory proteins, including succinate dehydrogenase/fumarate reductase subunits, pyruvate dehydrogenase components, thioredoxin, and Fe-S assembly proteins, together with increased riboflavin pathway abundance and low residual riboflavin levels, indicate enhanced respiratory cofactor demand and active flavin metabolism during lactate growth. These observations are consistent with previous studies linking lactate metabolism and microaerobic respiration to efficient B12 production [2]. Notably, lactate cultures displayed the highest biomass-specific B12 accumulation despite lower biomass formation, indicating that efficient corrinoid biosynthesis is associated with a balanced respiratory metabolic mode rather than maximal carbon throughput.

### Aspartate links redox balancing with B12 precursor allocation

Our data further demonstrate that carbon source-dependent metabolic adaptation strongly influences allocation of aspartate between redox balancing and corrinoid biosynthesis. Aspartate supplementation increased volumetric and specific B12 production during lactate metabolism but reduced B12 yields under fructose conditions. These opposing responses indicate that under lactate metabolism aspartate primarily supports biosynthetic precursor supply, whereas during hexose metabolism it is preferentially redirected toward succinate-forming pathways that support cellular redox homeostasis. Thus, metabolic competition between redox homeostasis and corrinoid precursor availability appears to be a key determinant of B12 biosynthetic efficiency.

### Corrinoid biosynthesis reflects metabolic state rather than enzyme abundance alone

The proteomic patterns observed for corrinoid biosynthesis further support this interpretation. Hexose metabolism, particularly fructose growth, enriched early-stage B12 biosynthesis enzymes and cobalt transport systems, consistent with high volumetric B12 production and elevated trace metal demand during carbohydrate-driven metabolism. In contrast, lactate preferentially enriched riboflavin biosynthesis and flavin-associated pathways, supporting respiratory metabolism and efficient biomass-specific corrinoid accumulation. The condition-specific regulation of the terminal enzyme BluB-CobT2 relative to upstream corrin ring biosynthesis proteins further suggests that final cobalamin assembly may be partially decoupled from precursor synthesis. Together, these observations indicate that B12 biosynthesis emerges from systems-level interactions among precursor availability, respiratory activity, redox balance, and carbon flux distribution rather than from biosynthetic enzyme abundance alone.

### Implications for metabolic state transitions and B12 production strategies

The integrated model proposed here **(Figure 8)** describes two alternative metabolic configurations in PFR. Lactate supports a respiratory, flavin-associated, and biosynthetically efficient state favoring high cellular B12 accumulation, whereas glucose and fructose establish a high-flux physiological configuration requiring enhanced pentose phosphate pathway activity and alternative electron-sink pathways to maintain redox homeostasis. Because substrates were supplied at equal mass concentrations rather than carbon-equivalent concentrations, the observed differences reflect both substrate identity and the distinct carbon and redox inputs associated with each compound. These findings expand current understanding of metabolic program transitions in PFR and highlight how carbon source determines the balance between growth, redox adaptation, and cofactor biosynthesis. From an applied perspective, the results support two-stage cultivation strategies separating biomass production from optimized B12 synthesis, for example by combining rapid biomass generation under hexose-rich conditions with subsequent metabolic transition toward microaerobic, respiratory states favoring efficient corrinoid accumulation.

## CONCLUSIONS

This study demonstrates that carbon source is a major determinant of metabolic organization in PFR, shaping how cells balance redox homeostasis, central carbon metabolism, and B12 biosynthesis. Integrated physiological, metabolite, amino acid, and proteomic analyses revealed that lactate and hexoses establish distinct metabolic configurations with different biosynthetic and redox-balancing priorities. Lactate supported a respiratory, comparatively redox-balanced state associated with amino acid biosynthesis, flavin metabolism, Fe-S cluster assembly, and efficient biomass-specific B12 accumulation. In contrast, glucose and fructose induced a high-flux, redox-constrained state characterized by increased carbohydrate turnover, pentose phosphate pathway activity, overflow metabolism, and activation of the aspartate–fumarate–succinate pathway as an alternative electron sink. The close association between aspartate metabolism, succinate formation, and aerobic viability suggests that aspartate-dependent redox balancing is an important adaptive mechanism during hexose metabolism. Our findings further indicate that B12 biosynthesis is regulated at the systems level rather than solely by biosynthetic enzyme abundance. Carbon source-dependent differences in corrinoid biosynthesis proteins, cobalt transport, riboflavin metabolism, and aspartate utilization suggest that precursor allocation, respiratory activity, and redox homeostasis collectively determine corrinoid biosynthetic efficiency. Together, these findings establish a mechanistic framework linking carbon source-dependent metabolic configurations with redox homeostasis and B12 biosynthesis in PFR, providing a basis for the rational optimization of industrial B12 production.

## Data availability statement

The proteomics data sets generated during and/or analysed during the current study are available in the PRIDE repository with dataset identifiers PXD079229. The other data sets generated during and/or analysed during the current study and not included in the manuscript are available from the corresponding author on reasonable request.

## Acknowledgements

Funding for this study was provided by the University of Helsinki and by Academy of Finland (project number 325784). Open access was funded by Helsinki University Library.

Mass spectrometry-based proteomic analyses were performed by the Proteomics Core Facility, Department of Immunology, University of Oslo/Oslo University Hospital, which is supported by the Core Facilities program of the South-Eastern Norway Regional Health Authority. This core facility is also a member of the National Network of Advanced Proteomics Infrastructure (NAPI), which is funded by the Research Council of Norway INFRASTRUKTUR-program (project number: 295910).

## Author contributions

KS: Formal analysis, Investigation, Methodology, Visualization, Writing – original draft, Writing – review & editing

BC: Formal analysis, Investigation, Methodology, Writing – original draft, Writing – review & editing

ME: Formal analysis, Investigation, Methodology, Visualization, Writing – original draft, Writing – review & editing

KH: Formal analysis, Investigation

PD: Investigation, Supervision, Writing – original draft SK: Methodology, Supervision, Writing – review & editing

TN: Formal analysis, Investigation, Methodology, Supervision, Writing – original draft, Writing – review & editing

VP: Conceptualization, Funding acquisition, Methodology, Resources, Supervision, Writing – review & editing

PV: Conceptualization, Funding acquisition, Project administration, Resources, Supervision, Writing – review & editing

## Conflict of interest statement

The corresponding authors confirm on behalf of all authors that there have been no involvement that might raise the question of bias in the work reported or in the conclusions, implications or opinions stated.

## Supplementary Materials

**Table S1.** MaxOutput file listing of all identified proteins from *P. freudenreichii* cells cultured on lactate (LAC), glucose (GLC) and fructose (FRC) as the carbon sources. Both the RAW and LFQ intensity values together with the identification type and the sequences of the matching peptides are shown. Proteins specifically identified from LAC-, GLC- and FRC-associated samples (detected in at least three out of five replica samples) are highlighted with green letters.

**Table S2.** Differentially abundant proteins identified from PFR cells across LAC, GLC, and FRC conditions. Normalized abundance values represent relative protein abundances in each condition; positive values indicate relative enrichment and negative values indicate relative reduction. ANOVA significance (“+”) indicates proteins differing significantly among conditions, while “Sign. pairs” lists significant post-hoc pairwise comparisons. Heatmap order and cluster color indicate hierarchical clustering-based grouping. Peptide counts, sequence coverage, molecular weight, score, intensity, MS/MS count, p-value/q-value metrics, protein IDs, and protein sequences describe protein identification and quantification quality.

**Figure S1.** Venn diagrams comparing the number of all identified proteins within five biological replica cells grown on fructose (FRC), glucose (GLC) and lactate (LAC). Proteome coverage was high in each three analysis; number of uniquely detected proteins was low in each 5 replica samples and over 96-99% of the identified proteins could be detected in 3 or more replica samples.

